# Measuring of the Quenching Rate of Atto-655 by Tryptophan

**DOI:** 10.64898/2026.04.27.721120

**Authors:** Niall G. Martin, Kasun Gamage, Lisa J. Lapidus

**Affiliations:** Department of Physics and Astronomy, Michigan State University, East Lansing, MI 48823

## Abstract

Intramolecular diffusion is an important, but often overlooked, property of intrinsically disordered proteins and plays an important role in folding, assembly and aggregation. Fluorescence resonance energy transfer (FRET) is used to observe reconfiguration over nanometer length scales while close range quenching over Angstrom length scales provides a complimentary view with different dynamics. There are several probe/quencher pairs that have been employed with varying levels of quantification of the quenching rate. Here we measure the electron transfer quenching parameters of the fluorophore Atto-655 by tryptophan using fluorescence correlation spectroscopy. Measurements with varying concentrations of quencher with low diffusion yield a distance dependent quenching rate. These parameters provide for a more quantitative analysis of measurements of intramolecular diffusion, particularly in crowded environments.

## Introduction

Over the past 25 years, the technique of measuring intramolecular diffusion in disordered proteins has been used to understand the basic polymer characteristics of polypeptides, early steps in protein folding, the relation between polymer dynamics and aggregation, and the behavior of proteins in crowded environments. The general concept of the measurement is shown in Fig. 1a. A polypeptide containing a probe and quencher is excited to a long-lived state and the state is monitored while the polymer reconfigures in equilibrium. If the quencher and probe come into close contact, the probe is irreversibly quenched back to its ground state, ending the measurement. This two-step kinetic model (see Fig. 1a) has a simple solution for the observed rate of the probe

**Figure 1.**
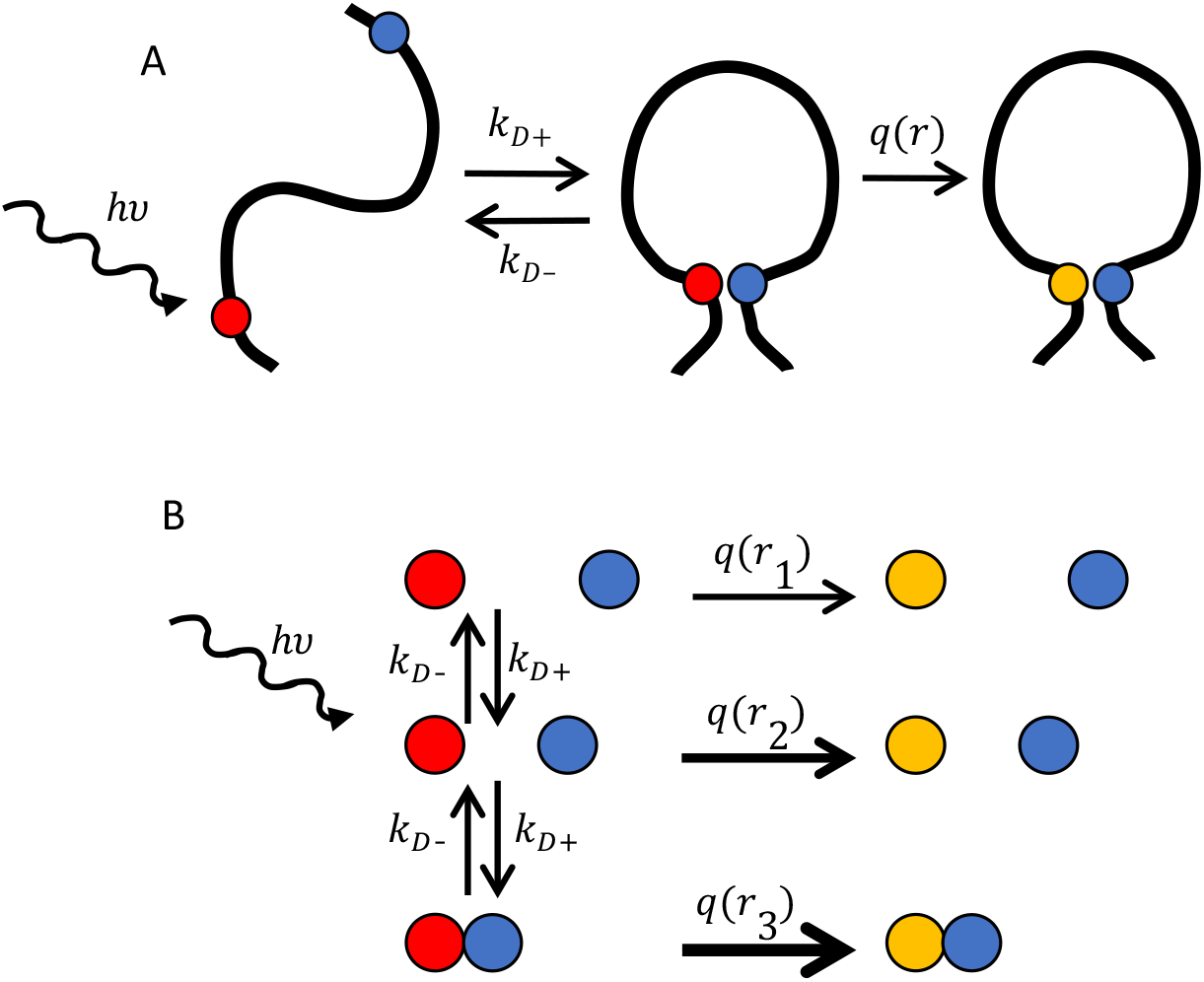
Schemes to measure intramolecular and intermolecular diffusion. A) A probe is excited to a long-lived state (red ball) that undergoes intramolecular diffusion in equilibrium. If the probe comes into contact with a quencher (blue ball), the excited state is irreversibly quenched (yellow ball), and the measured lifetime is given by Eq. 1. B) Bimolecular quenching at a variety of quenching rates depending on distance. Under crowding conditions, a range of quenching rates can be observed simultaneously.

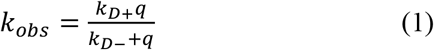

where *k*_*D+*_ and *k*_*D-*_ are the forward and backward diffusion rates and *q* is the quenching rate of the probe. A number probe/quencher systems were developed and employed to look at disordered peptides and proteins: 1) thioxanthone and naphtheline^1^, 2) DBO (2,3-diazabicyclo[2,2,2]oct-2-ene) and tryptophan^2^, 3) tryptophan and cysteine (Trp-Cys)^3^, and 4) Atto-655 and tryptophan (Atto-Trp)^4^, which all demonstrated close-range (< 1 nm) quenching. Additionally various FRET pairs of fluorophores have been used with a similar scheme but longer-range energy transfer^5^. While methods 1 and 2 require the attachment of one or both moieties to a polypeptide chain using solid-phase synthesis, method 3 requires no external labeling and can be used with recombinant expressed proteins engineered to have one Trp and one Cys in the sequence.

Method 4 uses a commercially available fluorophore that can be readily attached to Cys via maleiamide chemistry and can therefore be used with the same protein sequence as method 3.

Trp-Cys quenching (method 3) has been employed by our group for many years to measure a variety of sequences. The quenching rate, *q*, was determined to be due to electron transfer and therefore have a distance dependent quenching rate of the form

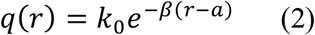

where *a* is the distance of closest approach, *k*_*0*_ is the rate at *a* and *β* is the decay distance. The distance dependent quenching rate implies that a variety of quenching rates can be observed as quenchers move towards and away from the probe for sufficiently high quencher concentration and sufficiently low diffusion, as shown in Fig. 1b. For the Trp-Cys system this was accomplished by embedding free Trp and Cys in a trehalose glass with Cys concentrations up to 1 M^6^.

While Trp-Cys quenching has proven a useful tool to measure many different sequences, it has some drawbacks. The probe is the triplet state of Trp which is observed by transient absorption of light at 450 nm. This signal is relatively weak and therefore requires a relatively large amount of sample at a moderate concentration and a long pathlength (typically 30 μM in 1 cm). This makes measurement of highly aggregation-prone, turbid or difficult to make samples impractical.

In contrast, Atto-Trp quenching employs fluorescence correlation spectroscopy of Atto-655 to measure intramolecular quenching for times much less than the translational diffusion time. The measurement takes place on a confocal microscope near the surface of a slide and can use turbid samples. Furthermore, the method requires very low concentrations (1-10 nM) and volumes (~20 μL) and overcomes the limitations of Trp-Cys quenching described above. Previous work has shown that Atto-Trp quenching in peptides is diffusion-limited and close range, but the parameters of the quenching rate have not been quantitatively determined. In this work we parameterize the quenching rate for the first time so that this method can be employed with greater precision in a wide variety of systems. The parameters are determined in samples with a low concentration of free Atto-655 (~1 nM), a high concentration of free Trp (up to 40 mM) and low translational diffusion (50% polyethylene glycol). To confirm these parameters, we use a highly crowded system to observe intermolecular quenching and solve the diffusion equation to reproduce the observed experimental dynamics. Finally, we take advantage of the fact both Trp-Cys and Atto-Trp quenching can observe the same protein sequence to compare intramolecular quenching of a disordered protein.

## Methods

### Protein expression

Human α-synuclein Y39W/A69C was recombinantly expressed as described previously^7^. Briefly, α-synuclein expression plasmids were transformed into BL21(DE3) competent cells by heat shock (42 °C, 90 s), followed by recovery in SOC medium for 1 h at 37 °C with shaking at 225 rpm. Cells were plated on LB agar supplemented with ampicillin (0.1 mg/mL), and successful transformants were selected by antibiotic resistance.

Starter cultures (5 mL LB containing 0.1 mg/mL ampicillin) were grown for 6 h at 37 °C with shaking, then transferred into 2 L LB supplemented with ampicillin (0.1 mg/mL). Cultures were grown overnight for 16 h at 37 °C with shaking, after which protein expression was induced with 1 mM IPTG for an additional 6 h. Cells were harvested by centrifugation at approximately 4,000 × g for 20 min at 4 °C.

Cell pellets were resuspended in lysis buffer consisting of 50 mM sodium phosphate and 300 mM NaCl, pH 8.0. Unless otherwise noted, all subsequent purification steps were carried out on ice or at 4 °C. Protease inhibitor, lysozyme, DNase, and RNase were added, and the suspension was stirred for 20 min. Cells were then lysed by sonication for 6 min total using 15 s on/15 s off pulses at 35% amplitude, followed by an additional 30 min of stirring.

The lysate was heated at 100 °C for 30 min, with gentle mixing after 15 min, and centrifuged at 30,300 × g for 20 min at 4 °C to remove precipitated contaminants. Ammonium sulfate was added to the clarified supernatant to a final concentration of 361 mg/mL. The solution was stirred for at least 30 min, incubated for an additional 30 min, and centrifuged at 34,500 × g for 25 min. The resulting pellet was stored at −20 °C.

For purification, the pellet was resuspended in 20 mM sodium phosphate buffer, pH 8.0, and subjected to anion-exchange chromatography using a 5 mL HiTrap Q FF column (Cytiva).

Protein was eluted with a linear gradient from 0 to 1 M NaCl in 20 mM sodium phosphate buffer over 20 column volumes. Fractions containing α-synuclein were pooled and buffer-exchanged into 20 mM sodium phosphate buffer prior to further purification by size-exclusion chromatography on a HiPrep 16/60 Sephacryl S-200 HR column equilibrated in 20 mM sodium phosphate buffer. Fractions were collected into tubes containing 1 mM TCEP. Protein-containing fractions were pooled and concentrated to approximately 600 μM.

K10C/W43Y GB1 was purchased from Bio-Synthesis. T2Q GB1 was a generous gift from the Pielak lab.

### Fluorescent labeling

Single-cysteine variants, Y39W/A69C α-synuclein and K10C/W43Y GB1, were labeled with Atto 655 Maleimide (ATTO-TEC). Protein samples in 20 mM sodium phosphate buffer, pH 7.1, containing 0.5 mM TCEP were incubated with dye at a 15.5:1 dye:protein molar ratio overnight at 4 °C. Excess unreacted dye was removed by size-exclusion chromatography using three HiTrap Desalting Columns packed with Sephadex G-25 resin connected in series (Cytiva).

### Fluorescence Correlation Spectroscopy Instrument

FCS is performed on a confocal microscope in which the photons emitted from a small volume on a microscope slide are divided equally between two detectors. The arrival times in each detector is recorded separately and a histogram is created for the times between photons. A HeNe laser illuminates the back port of an Olympus IX51 inverted microscope and focused with a 1.2 NA objective (Olympus U PLAN S-APO 60X) ~20 μm above the surface of a 170 μm-thick coverslip. Light emitted from a fluorescent sample is collected by the objective, passed through a dichroic mirror (Chroma T747lpxr) and out the camera port. The light is then focused onto a 200 μm pinhole, collimated again and split 50/50 between 2 single photon avalanche detectors (PDM). The laser power at the sample is typically 180 μW. Using the measured translational diffusion time of free Atto-655 and the known diffusion coefficient (40.6 Å^2^/ns), the confocal volume is determined to be ~3-4 fL.

Photon arrival times from two single photon avalanche detectors (PDM) were recorded by the MultiHarp 150 4N (PicoQuant) and analyzed with custom Matlab programs, a generous gift from the Enderlein lab. Data was collected in 30 second increments to filter out rare instances of large particulates traversing the laser beam and all data sets were analyzed together with a time resolution of 50 or 100 ns.

To fit the intermolecular quenching data of free Atto-655 and free Trp using the scheme in Fig. 1b, we assume the correlation on short time scales is related to the survival probability of the excited state of Atto-655, *S*(*t*|*r*), as^6^

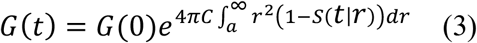

where *C* is the concentration of quenchers (in Å^-3^), and *a* = 4 Å is the distance of closest approach. We further assume an isotropic distribution of quenchers. The radially weighted function *F*(*t*|*r*) = *r*^2^*S*(*t*|*r*) obeys the diffusion equation

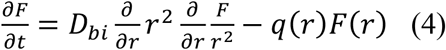

where *q(r)* is given in Eq. 2 and *D*_*bi*_ is the translational diffusion coefficient. To solve Eq. 4 we generate a rate matrix ***R***

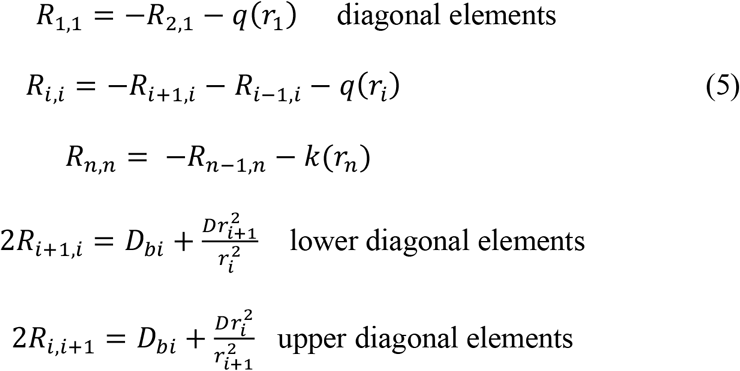

The matrix equation 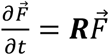is solved numerically with the boundary conditions that *F* is reflecting at *r = a* and *r = ∞* and *F*(0|*r*)=*r*^2^.

To fit the GB1 intermolecular quenching data, we follow the same procedure for Atto/Trp except Eq. 4 is modified to

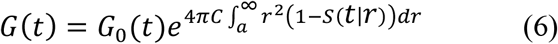

where *G*_*0*_*(t)* is the correlation due to translational diffusion given by

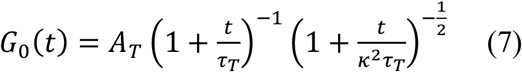

where *τ*_*T*_ is the time for the molecule to transit the confocal volume, *κ* is a geometric parameter that depends on the confocal volume and *A*_*T*_ = 1/*N* where *N* is the average number of molecules.

Intramolecular quenching FCS data were fit to the equation

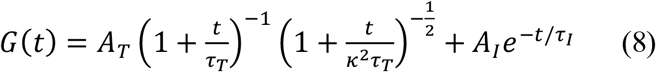

where *A*_*T*_, *A*_*I*_, *κ, τ*_*T*_ and *τ*_*I*_ are free parameters. *τ*_*I*_ =1/*k*_*obs*_is the decay due to intramolecular quenching.

### Trp-Cys Quenching

Trp-Cys quenching measurements were performed using pump-probe spectroscopy. A 266 nm UV pulsed laser beam (pump) with a 10 ns excitation and a 10 ms relaxation time is created from the fourth harmonic of an Nd:YAG laser (Continuum Surelite II-10) and converted to 289 nm by a 1 m Raman converter (LIGHT AGE) filled with 450 PSI of D2 gas. The produced 289 nm beam is used to excite the Trp to a long-lived triplet state. Then the lifetime of the Trp population in the triplet state was probed between each pulse by transient absorption using a continuous wave LASEVER 445 nm diode laser (probe). The beam intensity is recorded by a nanosecond photodetector (New Focus) and amplified by a LeCroy DA1855A differential amplifier. The output is then fed into a digital oscilloscope (Tektronix TDS 3032B) and the data is collected through a custom LabVIEW program. The temperature of the sample cuvette is a controlled by a Peltier sample holder (Quantum Northwest).

To prepare the sample for measurement, 2.7 mL of 50 mM Tris-HCl was degassed for 1 hour using N_2_O in a sealed long neck quartz cuvette (Hellma) to remove any oxygen molecules and scavenge free electrons that can quench the tryptophan triplet state or introduce photophysical effects to the measurements. TCEP at 10X the concentration of the protein was added to reduce any disulfide bonds. After degassing, the 300 μL of 300 μM α-synuclein was injected to reach a final concentration of 50 μM. The sample is then further deoxygenated for ~10 min by allowing N_2_O was allowed to flow on liquid surface.

To analyze the data, Eq. 1 can be rewritten as

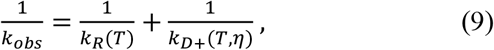

where

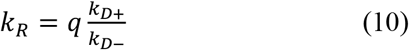

is the reaction-limited rate and *k*_*D+*_ is the diffusion-limited rate. These rates are determined by varying the viscosity (*η*) of the solution at a constant temperature (22° C). A plot of 1/*k*_*obs*_ vs. *η* is fit to a line in which the intercept is 1/*k*_*R*_ and the slope is 1/*ηk*_*D+*_.

### Szabo, Schulten and Schulten (SSS) theory

Quantitative analysis of intramolecular dynamics employs the use of Szabo, Schulten and Schulten theory, which models polymer dynamics as diffusion on a 1-dimensional potential. Using SSS theory, the reaction-limited and diffusion-limited rates defined in Eq. 2 are given by^8^

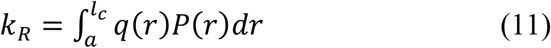

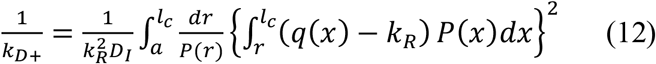

where *a* = 4Å is the van der Waals contact distance, *l*_*c*_ is the contour length of the chain, *r* is the distance between the Trp and the Cys, and *D*_*I*_ is the intramolecular diffusion coefficient.

### COCOMO simulations

Coarse grained simulations of α-synuclein at 50µM were carried out using COCOMO2.^9^ Simulations were performed using OpenMM 7.7.0. Langevin dynamics was applied with a friction coefficient of 0.01 ps−1, with temperature fixed at 298 K. The initial conformation for a protein chain was randomly assigned such that the chain doesn’t overlap with itself using a custom python script. Then each protein chain was placed randomly in a 100 nm size box such that no residue is within 1 nm of each other until the required concentration is reached.

Simulations were run for 4 µs at time steps of 0.02 ps saving every 20 ps. The first 200 ns of the simulation was considered as system equilibration and disregarded. ~ 90% of the conformations in the systems were monomeric, and multimers were rejected in analysis. Distances between the Trp and the Cys of the same chain were calculated for all molecules in the system for the complete run and binned by 0.1 Å into a histogram to obtain *P(r)*.

## Results and Discussion

To parameterize the distance-dependent quenching rate of Atto-Trp, we require a system of quenchers that have sufficiently high concentration and slow diffusion that different quenching rates at various distances can be detected. To achieve low diffusion, we use 50% polyethylene glycol (PEG) 8K which has a viscosity 1-2 orders of magnitude greater than water. The translational diffusion time of Atto in water and 50% PEG differ by 32x, in agreement with the viscosity estimate, as shown in Fig. 2a.

**Figure 2.**
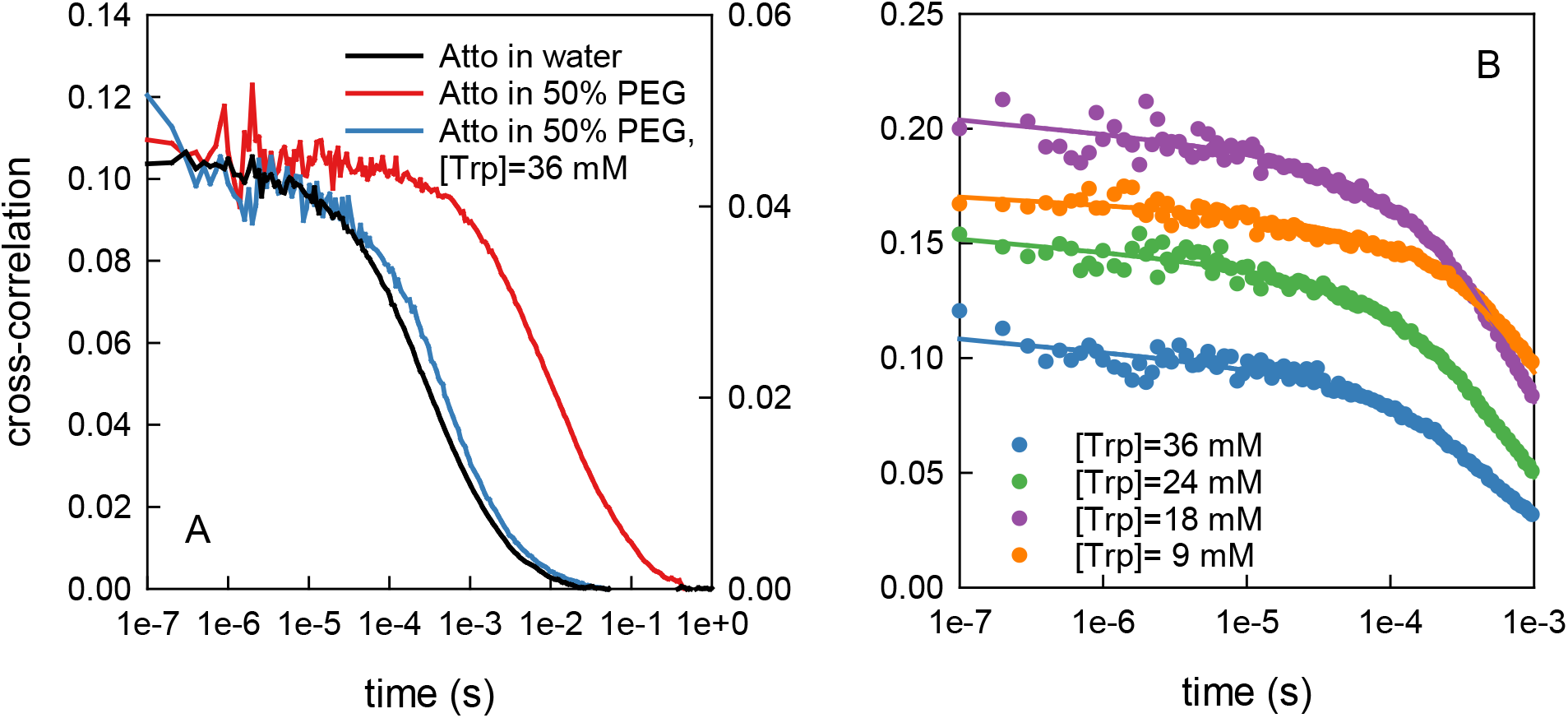
Tryptophan quenching of Atto-655 in 50% PEG. A) Comparison of cross correlation of Atto-655 fluorescence in water (right axis) and 50% PEG (left axis). B) Cross correlation of Atto with various concentration of Trp in 50% PEG (points). The lines are a global fit to all data for *t* < 1 ms, the time range over which the correlation without Trp is constant. Variations in amplitude in both plots are due to slight differences in the Atto concentration (1-5 nM).

To achieve high concentration, we use zwitterionic tryptophan which has a solubility limit of ~11 mg/mL in water and we were able to achieve 8 mg/mL in 50% PEG. Compared to the measurement in PEG without Trp, the primary decay is much shorter, indicating that this represents quenching, not translational diffusion (see Fig. 2a). Fig. 2b shows correlations of Atto in 50% PEG with various concentrations of Trp. The data were fit simultaneously to Eq. 3 with *k*_*0*_, *β* and *D* as global parameters and concentration, amplitude and offset as individual parameters for each data set. The global fit yielded a surprisingly low translational diffusion coefficient, *D*_*T*_ = 3.9×10^-10^ cm^2^s^1^, about 300x slower than would be expected from the translational diffusion in 50% PEG. This suggests that diffusion is anomalously slow over the short distances and sub-millisecond times over which quenching is observed. The fitted quenching parameters are given in Table 1. Fits with a range of starting conditions suggest that *β* is constrained to between 0.8 and 1.3 Å^-1^ while *k*_*0*_ much less constrained to greater than 4×10^9^ s^-1^. The small value of *β* compared to Trp-Cys quenching means that the quenching rate is significant over a longer range of distances. To confirm these quenching parameters, we look at Atto-Trp quenching in 2 protein systems.

| | $D_{bi}$ (cm<sup>2</sup>s<sup>-1</sup>) | $k_o$ (s<sup>-1</sup>) | $\beta$ (Å<sup>-1</sup>) |
| --- | --- | --- | --- |
| Atto/Trp in 50% PEG | $3.9 \times 10^{-10}$ | $1.2 \times 10^{10}$ | 1.2 |
| Atto/Trp in GB1 | $3.7 \times 10^{-11}$ | $1.4 \times 10^{10}$ | 0.96 |
| Trp/Cys in trehalose<sup>6</sup> | $1.5 \times 10^{-13}$ | $4.2 \times 10^9$ | 4.0 |

### Bimolecular quenching in a crowded protein system

The B1 domain of protein G (GB1) is an extremely stable protein that can be dissolved at concentrations as high as 100 mg/mL and remain folded. The wildtype sequence has one Trp at position 43 and no cysteines. We used a mutant, W43Y K10C to remove the Trp and add a Cys for Atto-655 labeling and dissolved this at low concentrations with 90 mg/mL of the wildtype. Fig. 3 shows the fluorescence correlation data has two decays, the slower decay is due to translational diffusion and the faster, non-exponential, decay is due to bimolecular Atto-Trp quenching. The data was fit to Eq. 6, yielding slightly different values for *k*_*0*_ and *β* than obtained above (see Table 1), but the data is adequately fit by those values as well, as shown in Fig. 3. *D*_*bi*_ is 10x slower than in Atto-Trp in 50% PEG and 4 orders of magnitude lower than measured for translational diffusion from the measurement of *τ*_*T*_. This may indicate anomalous diffusion of a highly crowded system or a slow structural change in the protein that is required to get the Atto and Trp within ~10 Å.

**Figure 3.**
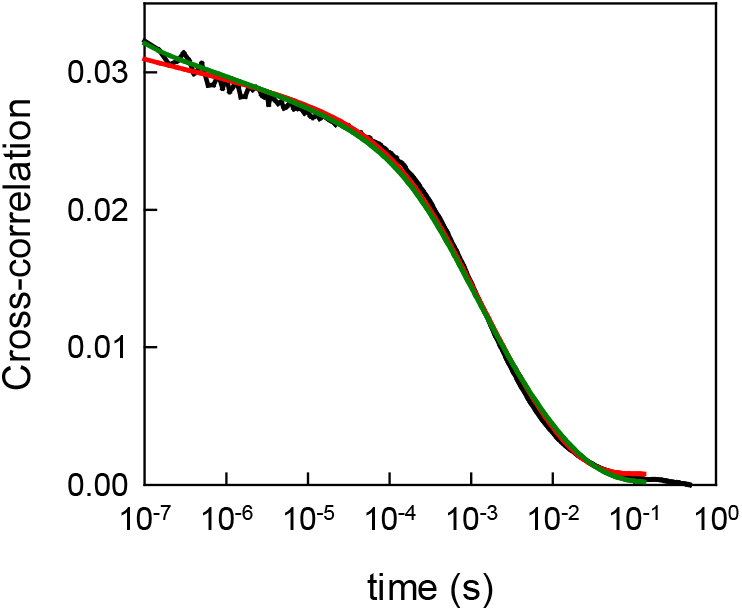
Atto-Trp quenching in crowding conditions. Fluorescence correlation (black) of 15 nM Atto655-labeled GB1in 90 mg/mL of unlabeled GB1. The red line is the fit to all parameters in Eq. 3. The green line is the fit to concentration, diffusion coefficient, amplitude and offset and fixing *k*_*0*_ = 1.2×10^10^ s^-1^ and *β* = 1.2 Å^-1^, which yields, *D*_*bi*_= 6.4×10^-11^ cm^2^s^-1^, approximately 2x the value in the full fit.

### Comparison of quenching in a disordered protein

Finally, we compared intramolecular quenching by Trp-Cys and Atto-Trp in α-synuclein. Eq. 9 shows that the measured intramolecular lifetime by either method (*τ*_*I*_ = 1/*k*_*obs*_) can be separated into these two rates, and we assume that while both rates can depend on temperature, only the diffusion-limited rate depends on the viscosity of the solvent (*η*), which is controlled with the addition of sucrose. FCS of Atto-655-labeled protein yields 2 decays from intramolecular quenching on the microsecond timescale and translational diffusion on the millisecond timescale (see Fig. 4a). FCS measurements are fit to Eq. 8, and Trp-Cys measurements are fit to 2 exponentials, the faster of which represents *k*_*obs*_. Previous measurements of Trp-Cys quenching in disordered proteins have shown that observed dynamics are typically neither reaction-nor diffusion-limited but rather a combination of the two^8^. A plot of viscosity vs. observed lifetime by Trp-Cys quenching yields a line where the intercept is 1/*k*_*R*_ and the slope is 1/*ηk*_*D+*_, as shown in Fig. 4b. In contrast, measurements of Atto-Trp quenching follow a line with an intercept consistent with 1/*k*_*obs*_ = 0, indicating that the observed rate is diffusion-limited (*k*_*obs*_ = *k*_*D+*_).

**Figure 4.**
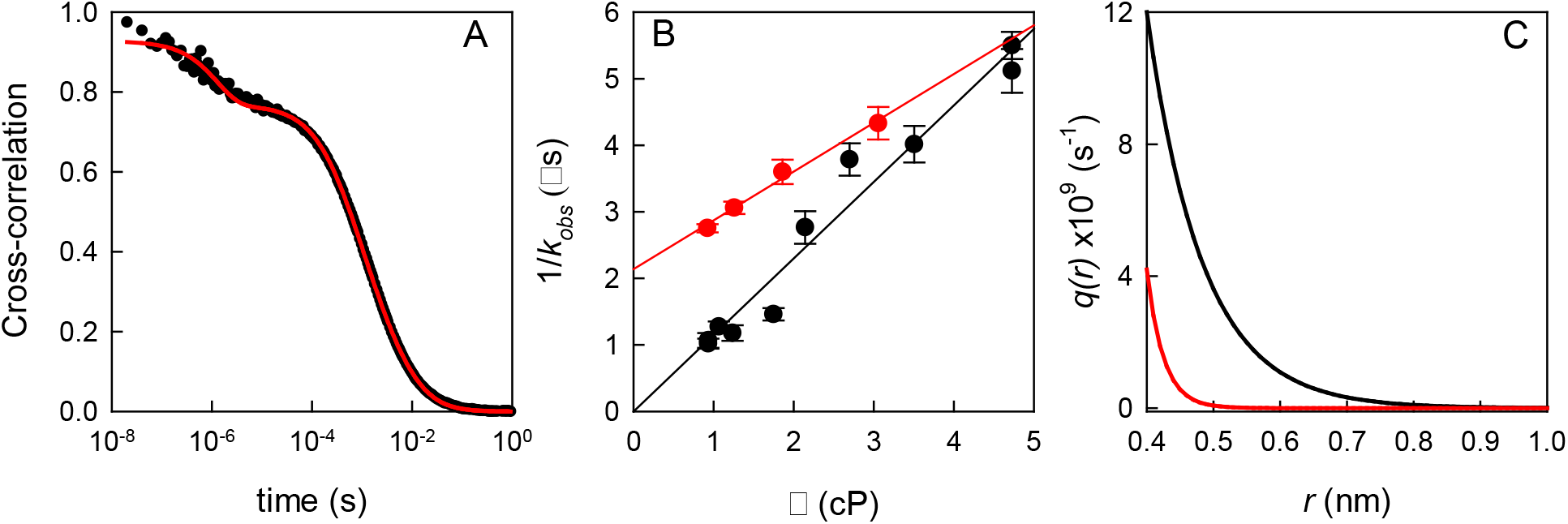
Intramolecular diffusion in α-synuclein. A) Fluorescence correlation data (black) of α-synuclein in 5% sucrose and fit to Eq. 6 (red). B) Intramolecular quenching rate by Atto-Trp (black) and Trp-Cys (red) quenching. The lines are linear fits 1/*k*_*obs*_ *=* (1.15±0.04)x10^-6^ η (black) and 1/*k*_*obs*_ = (0.73±0.06) x10^-6^ η + (2.14±0.11)x10^-6^ (red). C) Calculated quenching rates for Atto-Trp (black) and Trp-Cys (red) using the parameters in Table 1 and *a* = 0.4 nm

Measurable reaction-limited rates in Trp-Cys quenching allows the experiments to test computational assessments of the probability distribution in variety of sequences using Eq. 12, where *P(r)* is provided by simulation and the parameters of *q(r)* given in Table 1. We have previously shown that a coarse-grained model, COCOMO^9, 10^, produces probability distributions that accurately describe the conformational ensembles of disordered proteins. We carried out simulations using the improved version, COCOMO2 for a 50µM α-synuclein system and calculated *P(r)* as described in methods above. This probability distribution, when used in Eq. 12 with *a* = 5.45 Å, accurately reproduces the experimentally measured reaction-limited rate from Trp-Cys quenching. Assuming the same *P(r)* and *a* for Atto-Trp quenching, the parameters in Table 1 yields *k*_*R*_= 1.2×10^7^ s^-1^, significantly higher than *k*_*R*_ for Trp-Cys quenching and within the error of the intercept in Fig. 4. Using Eq. 13, we determine the intramolecular diffusion coefficient *D*_*I*_ = 5.1×10^-7^ cm^2^s^-1^ for Trp-Cys quenching and *D*_*I*_ = 2.5 x10^-7^ cm^2^s^-1^ for Atto-Trp quenching. This difference based on the probe/quencher pair likely reflects anomalous diffusion that depends on both length and time scales. Woodard, et al. compared Trp-Cys quenching with FRET for α-synuclein and found FRET to have faster intramolecular diffusion. MD simulations confirmed that the diffusion coefficient was distance dependent with longer-ranged probes giving higher *D*_*I*_.^11^

Compared to the Trp-Cys system, the Atto-Trp nominal quenching rate (*k*_*0*_) is significantly faster and the quenching distance (1/*β*) is significantly longer (see Fig. 4c). This leads to much higher rates of quenching over 2 Å beyond the distance of closest approach. While the fact that intramolecular Atto-Trp quenching is typically diffusion-limited prevents the testing of various probability distributions, it does allow direct measurement of *k*_*D+*_ without the introduction of viscogens, which allows the use of this quenching system in more complex solution conditions, such as within biomolecular condensates. The application of both quenching systems on the same protein sequence utilizes the best of both methods.

## Acknowledgements

We thank Michael Feig, Fabio Sterpone, Jörg Enderlein and Ingo Gregor for many helpful discussions. Protein samples of T2Q GB1 and the plasmid for α-synuclein was a generous gift from Gary Pielak group at the University of North Carolina. FCS analysis software was a generous gift from the Jörg Enderlein group at the University of Göttingen. This work was supported by the National Science Foundation MCB 2210228.

